# Titanium Nanoparticle Regulates Innate Immunity

**DOI:** 10.1101/2025.06.13.658543

**Authors:** Jacob J. Whittaker, David Eikrem, Joanna Whittaker, Gulaim A. Seisenbaeva, Karin Fromell, Kristina N. Ekdahl, Bo Nilsson, Mats Sandgren, Vadim G. Kessler

**Author notes:** Correspondence should be addressed to Jacob Whittaker. Authors contributed equally (Jacob J. Whittaker, David Eikrem).

## Abstract

*In-vivo* studies on wound healing have shown that small metal oxide nanoparticles (NPs), such as titania, ferria, and alumina, promote wound closure, likely by enhancing blood coagulation and via activation of the contact system. However, titania differs from the others as it supports the regrowth of normal skin tissue without scarring. This unique effect was hypothesised to result from its ability to suppress inflammation, possibly by interfering with the complement system. E*x-vivo* whole blood experiments within the present study confirm that titania NPs significantly dampen the immune response and surface plasmon resonance (SPR) biosensor analysis confirms that the nanoparticle binds with nanomolar affinity to complement C3, a central protein in the human innate immune system. To elucidate the molecular basis of this effect, we developed a novel cryo-electron microscopy (cryo-EM) methodology capable of resolving the challenging interface between metal NPs and proteins. This approach revealed that TiO□ binds with high specificity to a single domain on C3. The NP restricts the conformational dynamics of C3 and thus blocks the initiation of the complement cascade. These structural and biophysical insights support the biochemical findings and previous observations within the contact system. Our method advancement demonstrates that cryo-EM can uncover physiologically relevant interactions between proteins and nanomaterials at high resolution and offers a strong foundation for others investigating metal-biological interactions.

## Introduction

Engineered nanoparticles, such as silver, gold, zinc and titanium oxides, are products of modern science and have only been introduced into the environment within the last few decades. In particular, titanium dioxide (TiO_2_) nanoparticles have seen usage in an immense range of everyday products such as a whitening food additive (E171), confectionary, paint, hygiene and personal care products.^1-4^ Synthesised TiO nanoparticles can range from one nanometre in diameter through to several hundred nanometres, with each size class capable of eliciting distinct biological responses.^5^ Notably, macro-scale titania from implanted surfaces or environmental exposure can be gradually etched away by bodily fluids into smaller nanoparticle oxides, potentially contributing to systemic nanoparticle exposure.^6-8^ Despite the recent increase in anthropogenic metal-based nanoparticles and increased daily exposure, very limited safety information is available concerning their biological effect.^9^ Toxicity considerations are particularly relevant for TiO_2_ nanoparticles that are <5 nm in diameter due to their ability to cross the blood-brain barrier and penetrate cellular walls,^10, 11^ causing protein misfolding, enzymatic inhibition, dysregulation, aggregation and structural alteration.^12-15^ Previous work from our groups have unambiguously shown that a titanium nanoparticle 3.5 nm in diameter, in fact, improves burn wound healing and skin tissue regeneration in the contact activation system via upregulation (Figure 1).^16^

**Figure 1.**
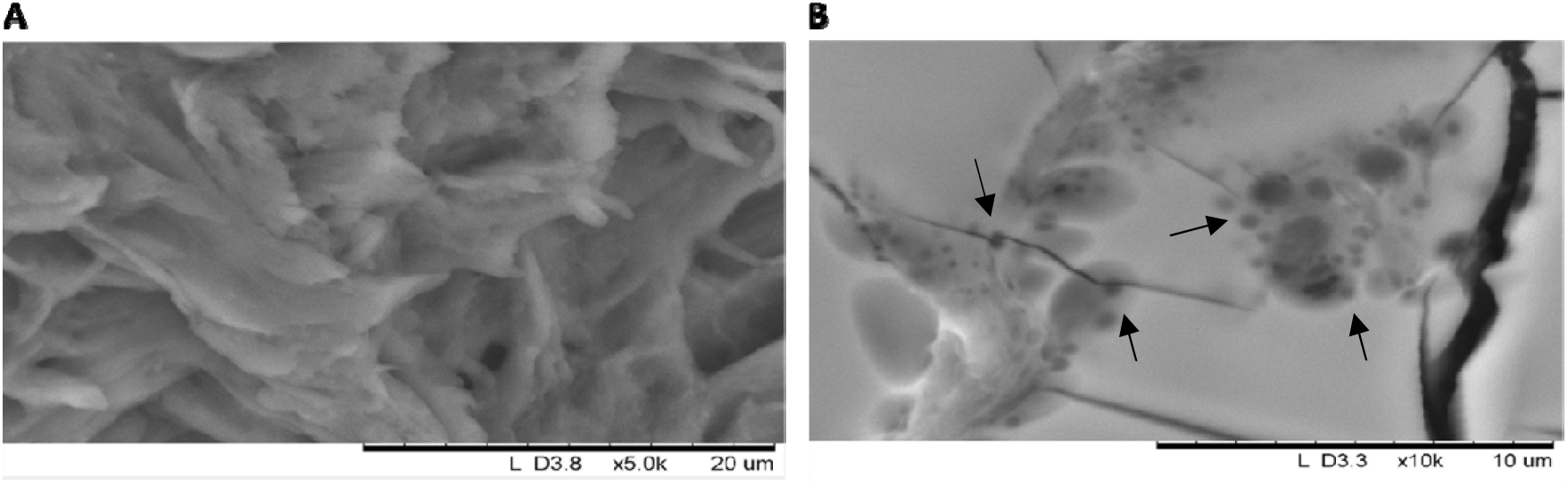
Comparative SEM images showing the influence that titanium nanoparticles have on human skin samples. A) Untreated human skin sample in the process of healing following trauma. B) Human skin sample coated with TiO_2_ nanoparticle.^16^

Throughout *in-vivo* studies in rats using these nanoparticles, we were able to enhance wound healing to such an extent that the regeneration of skin resulted in nearly no scar tissue. Using scanning electron microscopy (SEM), comparisons between naturally-healing skin and nanoparticlecoated human skin showed remarkable differences in the skin surface morphologies. Naturally-healing skin exhibited a randomly dispersed, fibrous surface, whereas skin coated with TiO_2_ appeared as smooth and well-organised with minimal evidence of trauma. Importantly, compared to the untreated control, there was a significantly quicker reduction of inflammation at the wound site. Other metal oxide nanoparticles were also tested, but none were able to elicit the same beneficial results as observed for titania. As the contact activation system operates together with other regulatory systems, we wanted to gain a molecular-level understanding into what was causing these nanoparticles to impart such considerable effects. This led us to begin investigating their influence on the complement system, a very closely-linked immune regulatory system.^17, 18^

Over the course of 500 million years, the complement system in vertebrates has evolved into the highly advanced and intricate defence mechanism found in humans today. This system involves the coordination of ∼50 different immune regulation factors and proteins.^19-21^ The complement system functions in tissue, plasma, intracellularly and demonstrates exceptional sensitivity to antibodies, foreign microbial sugars and pathogens.^22^ The C3 zymogen, an inactive enzyme precursor that requires activation to function, is perhaps the most significant protein in the entire complement cascade. It exists as the most abundant complement protein found in the blood of healthy humans at concentrations above 1.2 mg/mL.^23^ For activation of C3 to occur, a small 9 kDa domain (C3a) must be cleaved from the main 176 kDa chain (C3b) by a C3 convertase or serine proteases.^24-26^ This promotes a significant conformational shift of C3b and the exposure of a highly reactive thioester which is responsible for the opsonisation (tagging) of foreign particles for destruction by downstream immune system machinery.^27^ Importantly, dysregulation or dysfunction of C3 has been closely linked to diseases such as lupus and meningococcal meningitis, making it a clinically significant target for both understanding immune pathology and developing new treatments.^28-30^

Here we present the first biochemical and structural evidence of a physiological TiO_2_ nanoparticle downregulating the functional activity of wild-type human complement C3. Haemolysis results from whole blood models show that tiny quantities of TiO_2_ are enough to significantly decrease the activity of C3 and act as an alternative pathway-specific inhibitor. Furthermore, surface plasmon resonance (SPR) biosensor analysis shows that the NP has a remarkably strong affinity for C3. The K□value of 24 nM is six-times lower than that estimated for the original C3 commercial inhibitor, Compstatin Cp01.^31, 32^ Structurally, a major challenge in modern microscopy is the inability to visualise samples of mixed organic-inorganic species. We have overcome this obstacle and developed a methodology which has enabled us to observe this exciting interaction at viable resolutions for retaining molecular-level information. Our single-particle cryo-EM studies reveal a strong and specific interaction between a single TiO_2_ nanoparticle and protein domain is enough to deactivate the immune functionality of C3. The location of the bound nanoparticle restricts the motion of C3 and inhibits the formation of protein-protein interactions necessary for continued immune function. Our findings represent an important step towards enabling cryo-EM as a tool capable of capturing the communication between physiological proteins and interacting inorganic species. On one hand, we have provided valuable insight into how these physiologically-active TiO_2_ nanoparticles promote wound healing, while on the other hand, these effects are uncovered with molecular-level detail.

## Results

### Full-range of C3 flexibility revealed

The most compact state, shown in Figure 2a, appears as the dominant species in solution and represents the inactive state conformation. This arrangement shows the thioester domain (TED) compressed and adjacent to the macroglobulin ring (MG) domains of MG2 and MG6. The 120 Å diameter of this compressed structure supports the biophysical results and explains our observed discrepancy with previous structures (see ESI). The interactions between the TED and MG domains are facilitated mainly through dipole contacts rather than strong ionic interactions. This feature allows for the movement experienced by the MG domains to proceed freely. In fact, of the three interacting domains in this region, there are almost no electrostatic interactions or hydrogen bonds formed. Instead, a select few complementary hydrophobic and hydrophilic pockets of amino acids are enough to stabilise this conformation (Figure 3).

**Figure 2.**
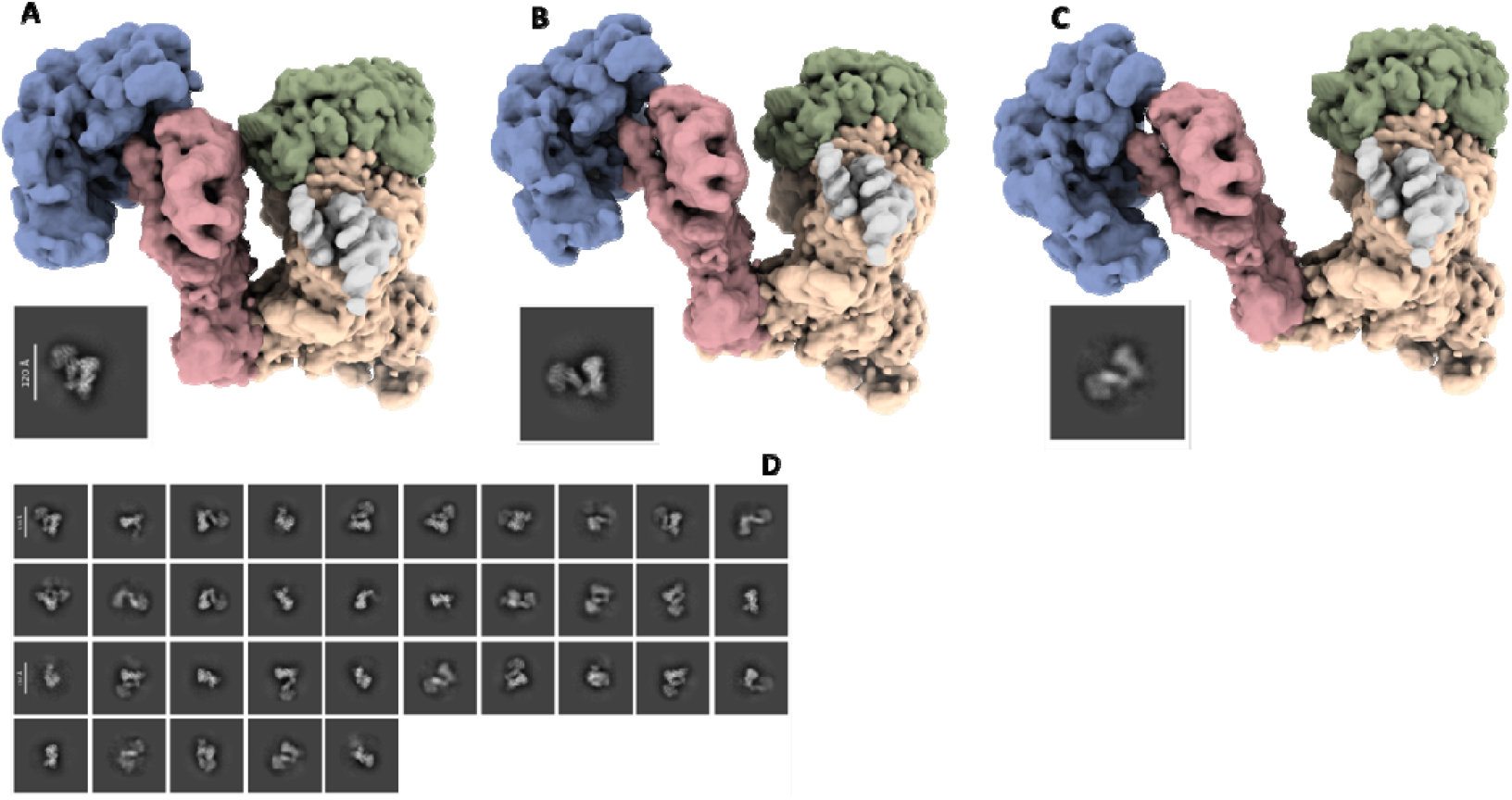
Cryo-EM densities and 2D classes showing the flexible heterogeneity of native C3 in a sample. The protein density is coloured to show the functional areas, such as C3a (grey), TED (green), connecting arm (red), MG head (blue) and body regions (beige). **A)** The most compressed, inactive conformation. **B)** An intermediate state exposing a gap between the central arm and TED regions. **C)** The extended state of C3 where the connecting arm region has moved away from the biologically-active TED, exposing it to potential foreign particles. **D)** 2D classes which exemplify the large conformational range achieved by C3.

**Figure 3.**
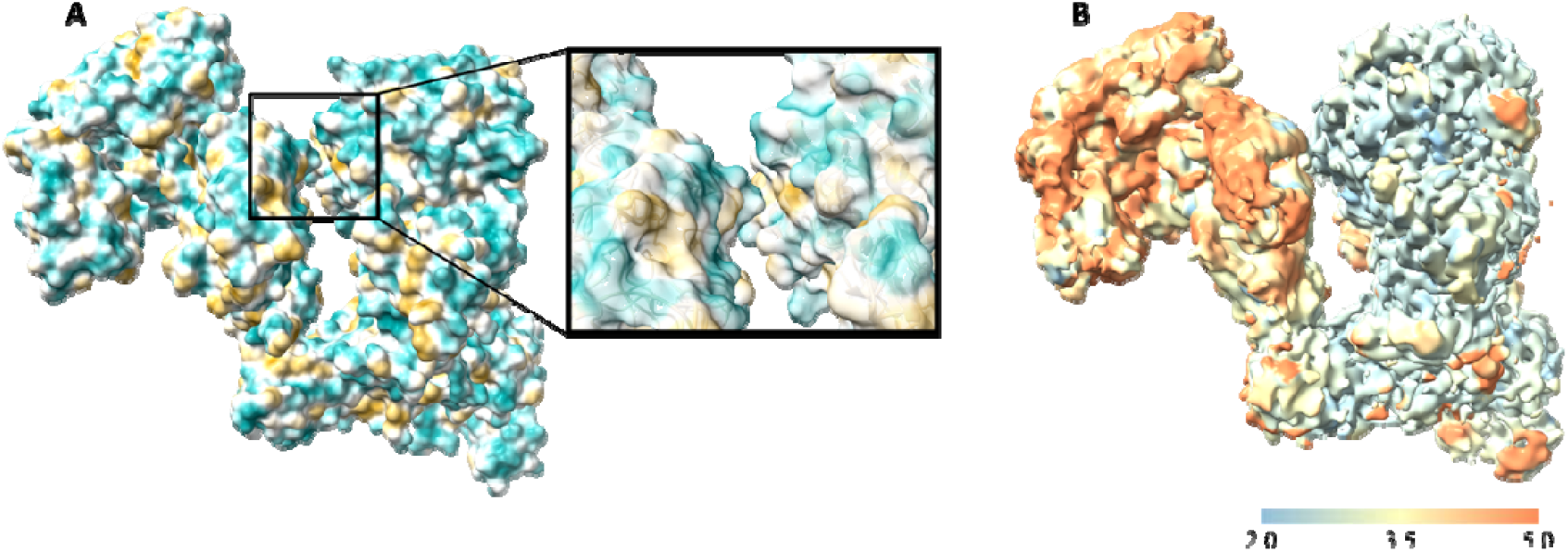
**A)** Native C3 electron density displayed as a hydrophobic (yellow) and hydrophilic (cyan) map with a close-up view of the interacting regions between the connecting arm and thioester domain. **B)** The local resolution electron map of native C3 calculated to FSC_0.143_.

3D Flexibility analysis shows two distinct states attributed to MG ring domains 1 - 6 which possess the ability to locally swivel and pivot in-place (ESI Figure S2). This was an unexpected finding, as the structurally well-characterised macroglobulin ring domain is reported as rigid.^33, 34^ Combining both 3D variability and 3D flexibility analyses, the global hinge-like motion between the contracted and expanded states could be resolved in addition to the flexible local motions experienced by each domain throughout the full movement. The MG domain movement between each conformational endpoint was found to reach up to 28 Å, with the MG2 and MG6 connecting arm region shifting by 12 Å. Minor classes found in 2D classification suggest that the extension may actually be greater than this, however these views were too rare to be reliably incorporated into the 3D variability constructions. In biological systems, the thioester domain acts as a “warhead” to connect to the detected foreign particle. The current structural series shows the MG2 and MG6 domains assist with this tagging process by uncovering the thioester region and exposing it for binding. These domains act as an extending arm which moves the bulky MG regions away from the TED by roughly 30°. Furthermore, the MG7 domain exhibits a slight bending motion which affords more solvent-accessible space for the linker domain to extend into.

Each “compact” and “stretched” conformation could be reconstructed to a FSC_0.143_ of 3.0 Å and 4.0 Å, respectively (ESI Supplementary Table 1). The local resolution range varied more depending on the degree of flexibility demonstrated in each conformational state. The alpha chain of C3 remained rigid with local resolution reconstructions of 2.5 Å possible. The flexible MG ring domains could be reconstructed to approximately 5 Å in the extended pose while a higher resolution of 3.2 Å was achievable in the contracted, more rigid conformation.

### TiO_2_ nanoparticle and human complement C3 interact with nanomolar affinity

Repeating the DLS and thermal unfolding experiments on TiO_2_ in the same buffered system as C3 proved unsuccessful as the presence of salt promoted the near-immediate aggregation and gelation of the nanoparticle colloid throughout a range of concentrations (ESI Figure S3a). Instead using only filtered water, the nanoparticles showed no signs of aggregation which was confirmed by DLS as the TiO_2_ particles were found to be of a homogenous diameter and stable at room temperature for 2 hours (ESI Figure S3b). After confirming that C3 was stable in filtered water, several concentrations of TiO_2_ nanoparticle were added to differing aliquots of 0.4 mg/mL C3 for DLS and nanoDSF analysis (ESI Figure S3c). DLS results showed no appreciable indications of protein or nanoparticle aggregation. NanoDSF showed near identical T_m_ inflection points for C3 in both water and buffered solution over a range of temperatures. Importantly, the addition of TiO_2_ resulted in an increase of T_m_ by ∼7 for several separate concentrations of nanoparticle, demonstrating a meaningful interaction and significant thermostabilising effect of the TiO_2_ species on C3.

Importantly, the interaction between the two species was confirmed and characterised via SPR biosensor analysis. The affinity between the nanoparticles and the C3 protein was high, as indicated by the estimated K_D_ of 24 nM (Figure 4). The poor fit using a 1:1 interaction model (Figure 4A) suggested a more complex binding event than a simple Langmuir interaction (i.e. a 1-step reversible interaction with a 1:1 stoichiometry). The improved fit using a two-state reaction (Figure 4B) supports an interaction involving a conformational change of the protein upon binding of the NP. Given the conformationally-heterogenous tendency of C3, it is unsurprising that the SPR biosensor data are consistent with ligand-induced conformational changes.^35^ This direct and time-resolved interaction mechanistic information therefore both supports the structural data, and reveals that the interaction has a higher affinity than Compstatin Cp01 (150 nM), the original commercial C3 inhibitor.

**Figure 4.**
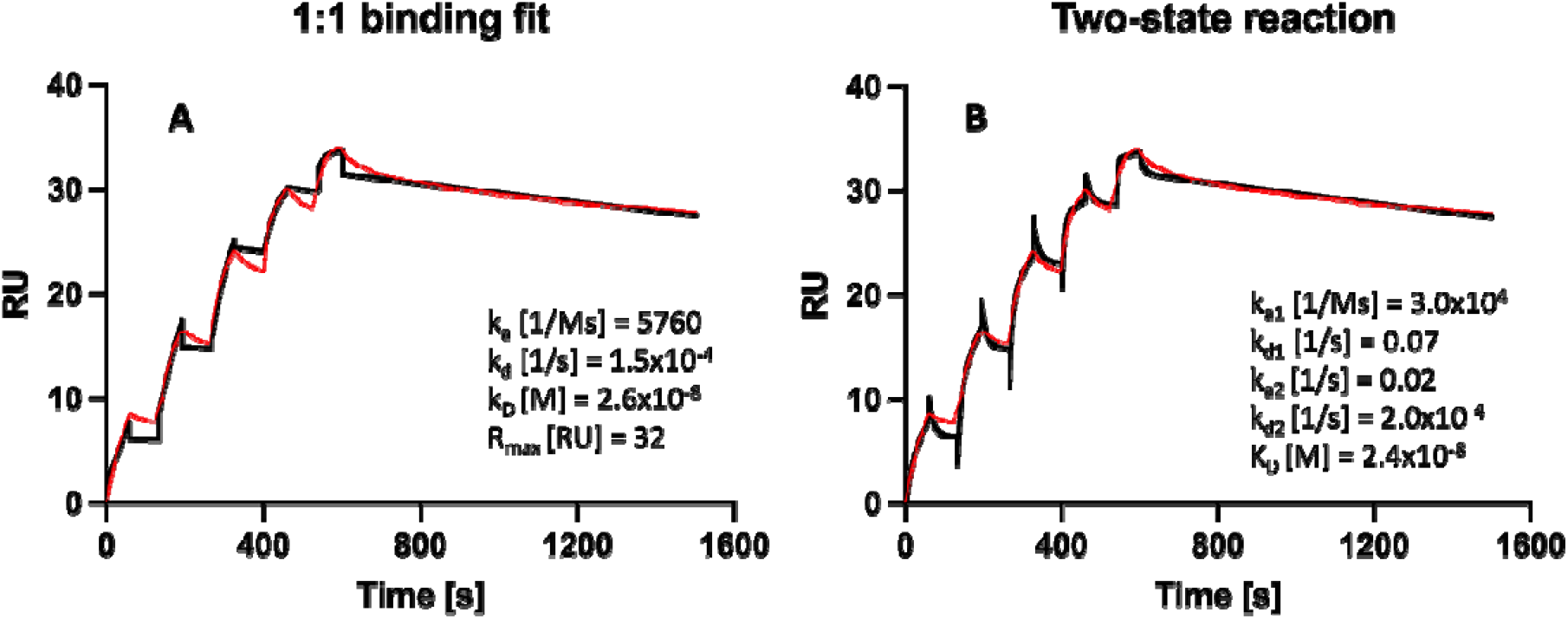
Sensorgrams of C3 interacting with immobilised TiO□ nanoparticles. Experimental traces from single cycle kinetic experiments are in red, theoretical traces from global, non-linear regression analysis fitting are shown in black. The sensorgrams represent a two-fold dilution series, starting from 10 µM down to 0.625 µM concentration. *A)* The theoretical fit and kinetic parameters obtained by a global non-linear regression using a 1:1 interaction model. *B)* The sensorgram fitting based on a two-state reaction model.

### Biochemical assays show TiO_2_ nanoparticle downregulates C3 activity

To test the effect TiO_2_ may have on C3 function, the formation of the alternative pathway (AP) convertase, C3bBb was considered. In the presence of Mg^2+^ ions, factor B (FB) binds to C3b forming the foundation of an AP convertase, allowing for the action of factor D (FD) to cleave FB into fragments Ba and Bb. In turn, C3bBb is formed with a substrate specificity for native C3, the convertase cleaves off C3a generating C3b. Using C3b as a positive control, native C3 (1 µM) was incubated with and without TiO_2_ (1 µM) prior to the addition of FB and FD, relying on C3(H_2_O) contamination or spontaneous hydrolysis occurrence to enable convertase formation and Bb generation. To analyse the effect of TiO_2_, the ability for FB binding to native C3 was assessed by comparing peak area values of FB and fragment Bb generation detected by a monoclonal antibody against Factor Bb neoepitopes. The capillary electrophoresis data generated combined area for Factor B and Factor Bb that were converted to percentage area in the Compass for Simple Western (v 6.1.0) programme after peak naming. Figure 5A conveys the percent of Factor B remaining, indicating a reduction in the AP convertase formation and activity, reducing Factor Bb generation. In the presence of TiO_2_, after 5 minutes, 95.1% FB remains (Native C3; 71.8%, C3b; 64.3%), 30 minutes 69.6% remains (Native C3; 13.9%, C3b; 15.6%) and 60 minutes 18.5% remains (Native C3; 11.9%, C3b; 8.9%).

**Figure 5.**
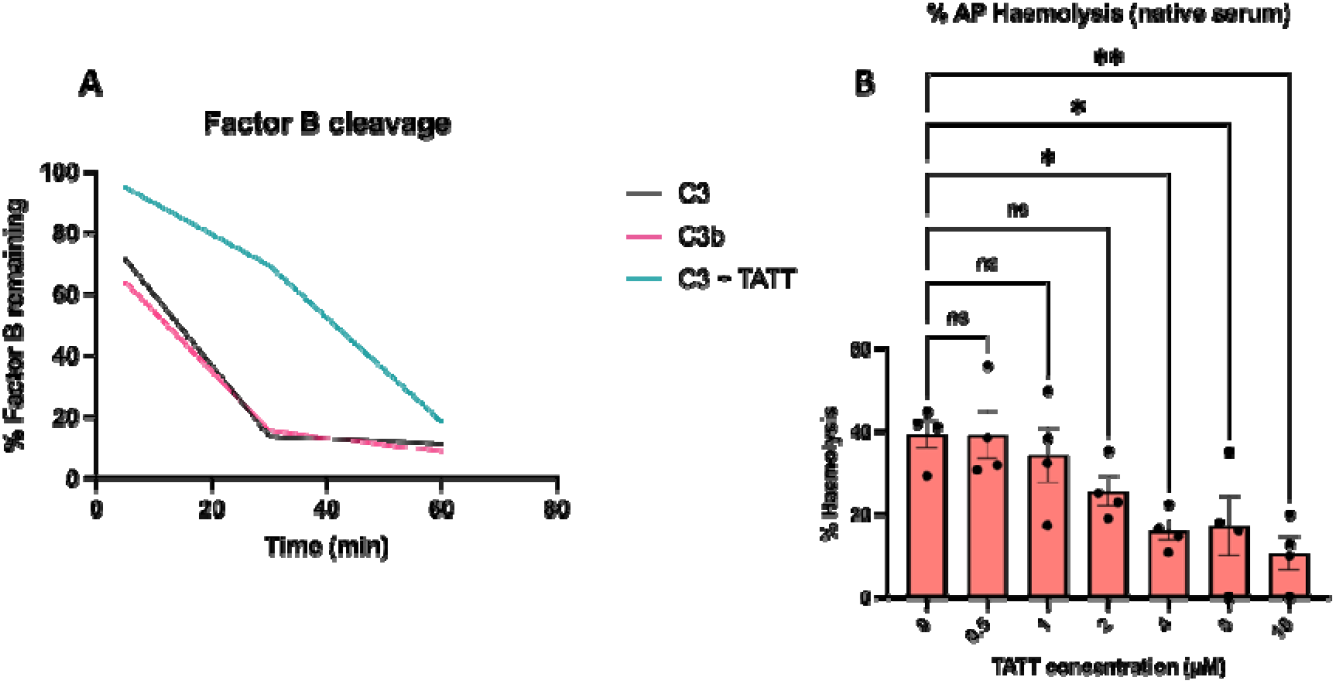
**A)** The percentage of factor B remaining after incubation of C3, C3b and C3 incubated with titania nanoparticles over time calculated from the representative Wes immunoassay. Percent calculated from peak areas across factor B and Bb populations in Compass for Simple Western software. **B)** Alternative pathway haemolysis assay of rabbit RBCs incubated with C3 and increasing TiO_2_ concentration. Measured at 540 nm and presented as an overall percentage of native serum. n=4, data presented as mean ± SEM, significance presented as \**P*< 0.0332 and *P*\*\*< 0.0021 determined by One-Way ANOVA with Dunnet multiple comparison.

### Haemolysis of red blood cells by C3 is inhibited with TiO_2_

Building on the AP inhibition, the capacity for native C3 to maintain its haemolytic properties was tested. Native C3 was incubated with incrementally increasing concentrations of TiO_2_ (0.5, 1, 2, 4, 8 and 16 µM). Rabbit red blood cells (RBCs) were used in this experiment as they fail to inhibit human complement and therefore become rapidly opsonised with C3b, enabling AP convertase and subsequent membrane attack complex formation (MAC) assembly, leading to haemolysis. In this serum dependant assay, an absorbance at 540 nm was recorded from native serum (total lysis), C3 depleted serum (blank) and the test samples of C3 with or without nanoparticles. The test samples are reported as a percentage of total haemolysis and show a significant reduction in haemolysis as nanoparticle concentration increases compared to neat native C3 (4 µM TiO_2_ *p*= 0.0187; 8 µM TiO_2_ *p*= 0.0268; 16µM TiO_2_ *p*= 0.003) as shown in Figure 5B.

### Direct observation of TiO_2_ nanoparticles interacting with human C3

Using these optimal conditions, C3 and TiO_2_ at differing ratios were applied to Quantifoil EM grids and plunge frozen in liquid ethane in preparation for cryo-EM. Despite their modest size of 3.5 nm and each nanoparticle possessing ∼1800 titanium atoms, the nanoparticles were readily visible as intense white spots on the micrograph with the 185 kDa protein in close proximity, barely visible as a blurred grey signal (ESI Figure S4a). Additional sample processing in RELION 5.0 provided more discernible particles of both C3 protein, TiO_2_ nanoparticles as well as classes which showed both in close proximity (ESI Figure S4b, S4d). Several ATLAS and eucentric height scans on different grids suggested that both particle distribution and contrast were favourable in thicker ice. It was immediately obvious from the micrographs that the nanoparticles further contributed to sample heterogeneity as they did not appear to interact with the protein in a uniform manner. This became evident from particle picking and 2D classification as several potential classes showed up to 3 nanoparticles may be interacting with a single protein, although at low alignment resolutions, this may be attributed to increased particle densities in the thick ice regions. Using a non-standard processing strategy in RELION 5.0 (see methods), we obtained a low-resolution electron density map suitable for protein and nanoparticle fitting.

Despite a loss of resolution, the C3 core particle density which comprised of the thioester and surrounding domains, matched very well with the previously-collected apo-C3 (amino acids ∼550-1641). Similarly to apo-C3, the macroglobulin ring density remained challenging to capture, however the location of the missing domains could be approximated as the MG2 and MG6 domain densities connecting the TED and MG ring, could be accounted for. The predominant pose of the C3 protein appears to be in the most compressed state, where the MG2 and MG6 domains rest directly parallel to the thioester domain. Several charged residues along the linker region form complementary interactions between oppositely charged residues on the external face of the thioester domain, further stabilising this conformation (Figure 6A, B). The vast majority of 2D classes also support this as we were unable to locate any rare views of extended protein in close proximity to nanoparticle density. In fact, it seems as though these extended forms of C3 are only observable in the absence of TiO_2_.

**Figure 6.**
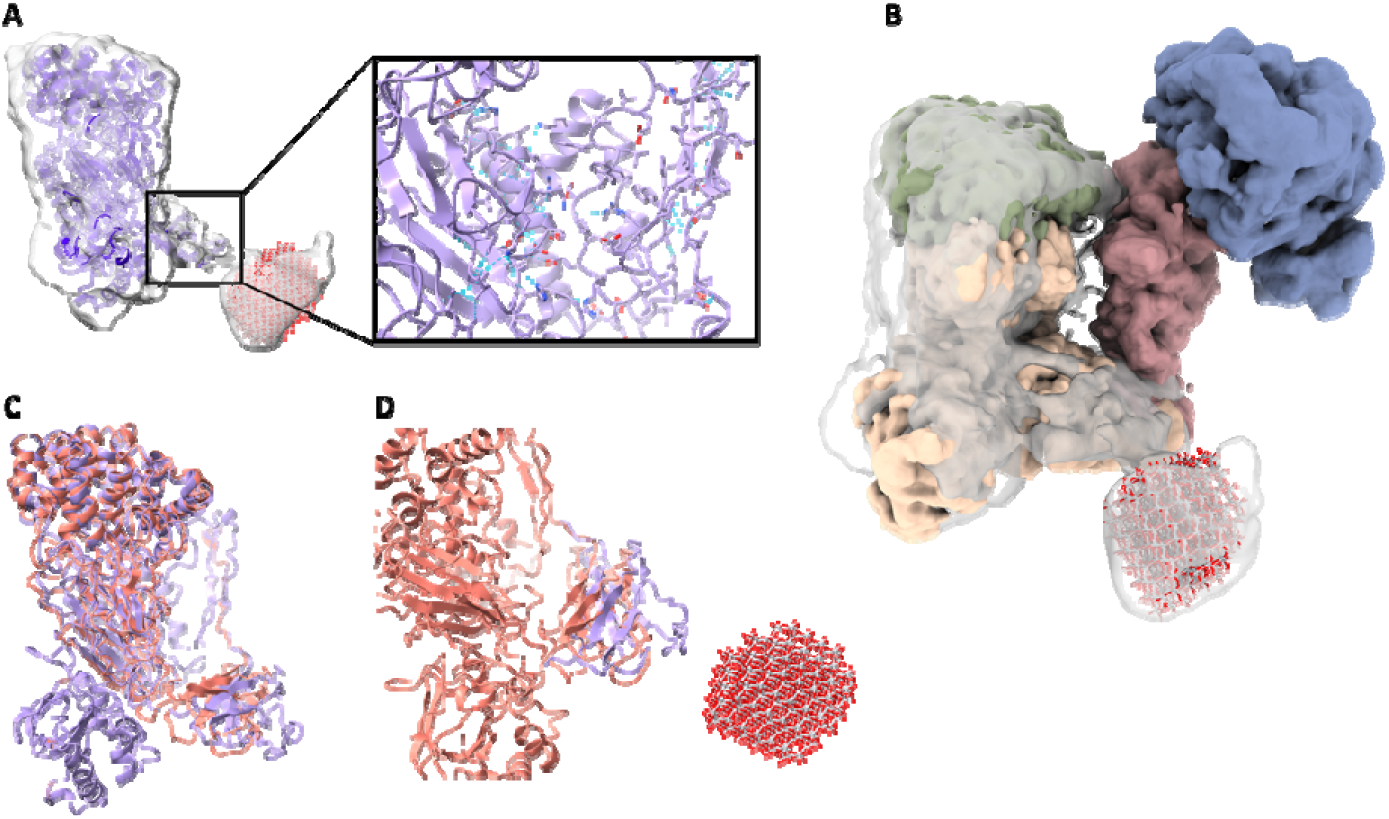
**A)** Electron density of the C3 core particle domains and the density attributed to the TiO_2_ nanoparticle. Top right: Intra-domain stabilising interactions between the linker and thioester domains. **B)** Overlaid electron density maps of the native C3 and TiO_2_-bound C3 structures highlighting the clear new density from the nanoparticle. **C)** Superimposed models of reported C3 (orange) and C3 from the current work (violet), demonstrating the disparity between the MG7 domain locations - RMSD 4.27 Å. *D)* Zoomed in image of the MG7 domains and the MG7 translated motion towards the TiO_2_ nanoparticle.

Although lacking a defined binding site, protein flexibility and alignment challenges from spurious classes, a clear density corresponding to the diameter of TiO_2_ was located adjacent to the C3 electron density. Owing to the signal thresholding during data processing, the most intense signal from the interior of the nanoparticle has been eliminated, leaving just the periphery which resulted in a density with a hollow interior. The pseudo-spherical density can be confidently attributed to the nanoparticle rather than the missing macroglobulin ring domains as the TiO_2_ density not only appears as the most intense after map contouring compared to protein signal, but is approximately 40 Å from the expected location of the macroglobulin ring. Furthermore, this region is situated well outside the latent axis of motion displayed by the apo-C3 structure. Overlaying the reported inactive structure with the nanoparticle-bound C3 shows that the majority of the structure is unchanged, however producing a calculated RMSD of 4.27 Å relative to the MG7 domains of the native and TiO_2_ structures (Figure 7A, B). This variability is largely due to the apparent outward shift of the MG7 domain (residues 807-911), as the RMSD of the remaining protein is 0.912 Å.

**Figure 7.**
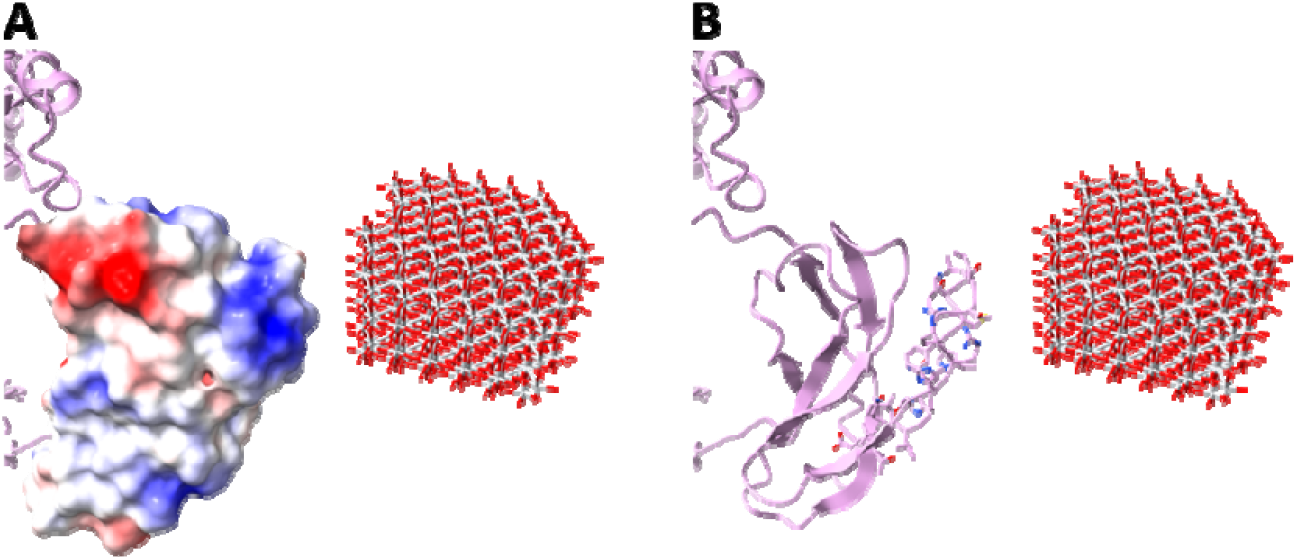
**A)** The electrostatic maps of MG7. Blue represents positively-charged domains, negatively-charged regions are shown as red. **B)** Stick representation of the interacting residues in MG7 with TiO_2_.

This macroglobulin domain is connected by two linker regions and is reported to move during activation of C3, demonstrating a rotation of approximately 35. This movement is facilitated by the excision of the polypeptide chain between MG7 and MG8 by factor I, thus allowing the two MG domains to nearly exchange positions.^36^ Within an inactive C3, this domain is situated between the linker and thioester domains, and has already been shown to move within the inactive state in the apo-C3 structure. The location of MG7 in the current structure shows an outward shift of approximately 9 Å in the opposing direction to MG8. As the linkers are still intact, a slight rotation of this domain is afforded to form an interaction with the nanoparticle which is positioned 6 Å adjacent to this domain. Given the significant electronegative nanoparticle, it was anticipated that any interaction formed between the species would occur at an acidic or phosphorylated region of the protein. A close inspection of the solvent-accessible residues on MG7 indeed confirm that the nanoparticle likely participates in an outer-sphere complexation with several electropositive residues (His846, Lys857, Arg858, Arg859, His860) in addition to hydrogen bonding interactions with several more residues (Cys851, Thr855, Thr856, Gln862) within the region.

## Discussion

### Physiological impact of titanium nanoparticles in the complement system

Titania surfaces have proven to be incredibly useful, particularly within medical devices and surgical implants, yet recent evidence shows that small (<10 nm) nanoparticles can be leached by the blood from these bulk surfaces. To the best of our knowledge, our work represents original insights into the physiological effects of small TiO_2_ nanoparticles on our immune system, following leaching from implanted surfaces. Significant improvements in the physical properties of complement C3 with TiO □ indicated direct communication between the species at a range of concentrations. This was surprising as neither of these components had any engineered binding sites specific for the other. Furthermore, such an interaction was unable to be repeated for the other complement proteins of C3a, factor B and factor D, in the alternative pathway, thus suggesting that there was an element of protein selectivity. Confirmation of C3-TiO_2_ specificity was quantified by SPR biosensor analysis. The two-state reaction model provided a better fit to the experimental data than the simple 1:1 interaction model, although both yielded essentially the same K □ value. The improved fit obtained with the two-state model indicates that the binding is not limited to a simple bimolecular interaction, but instead involves an additional step, consistent with a conformational change in C3. Such behaviour is typical in SPR analyses where the two-state model describes an initial reversible binding event, followed by a slower conformational rearrangement that stabilises the complex. Taken together, these findings suggest that the interaction between C3 and immobilised TiO□ nanoparticles is more complex than a straightforward 1:1 binding process. The estimated K □ represents the affinity for the interaction with consideration to the more complex interaction.

As TiO□ nanoparticles of this size, charge and composition are leached into the bloodstream, it was pertinent to investigate their physiological impact in *ex-vivo* models using red blood cells. Results from the factor B binding assay showed that in the presence of TiO□, the formation of the C3 convertase was greatly hindered and unable to activate nearby C3. Importantly, prior biophysical information showed no interaction between TiO□ and factor B, suggesting that factor B was not being influenced by the nanoparticles. In combination, these data strongly indicated that the interference was from an altered complement C3 protein. To assess this, we performed an alternative pathway haemolysis assay to show any differences in the ability of C3 to naturally lyse red blood cells in the presence of TiO_2_. After 60 minutes in the absence of nanoparticle, 40 % of RBCs were lysed compared to 10 % in the presence of 16 µM TiO_2_, corroborating that the activity of C3 was selectively reduced by the nanoparticles.

### Structural basis for complement inhibition by titanium nanoparticles

We lastly sought to understand the molecular-level influence that the titanium nanoparticle imparted on complement C3 and how this key protein was being inhibited. Despite its small size, visualising such a metal-dense nanoparticle in the presence of proteins had evident repercussions on particle alignment and 2D classification challenges.^37-39^ Following a non-standard processing strategy, a low-resolution reconstruction shows the interaction between C3 and a single TiO_2_ nanoparticle. Notably, although the C3-TiO_2_ data are low resolution, it is clear by the predominant protein conformation that C3 indeed remains in an inactive state. The compact density produced a diameter of approximately 120 Å, much more similar to the size of inactive C3 compared to activated structures of 160 Å.^36^ Despite the lack of a dedicated binding site for the nanoparticle, the location of TiO_2_ does not appear to be random as it is situated adjacent to an electropositive, solvent-accessible region of the MG7 domain. In the reported C3 convertase structure (2WIN),^40^ the MG7 domain and C-terminal domain of C345C form a junction where factor B is able to bind (Figure 8). In our structure, the titanium nanoparticle is located where factor B would normally interact with C3 to activate the zymogen and form the convertase, crucially, this interference at the MG7 domain explains why binding of factor B to C3 was inhibited, ultimately resulting in the reduction of C3 convertase, C3bBb. No data suggested an equilibrium or competitive binding between TiO_2_ and factor B to C3. This can be justified as the nanoparticle, albeit not much larger than factor B, is an extremely electronegative species as the titanium core is coated by oxygen. This results in a steric and electrostatic repulsive effect on the binding of factor B to C3 once TiO_2_ is bound, producing no interaction between the two species.

**Figure 8.**
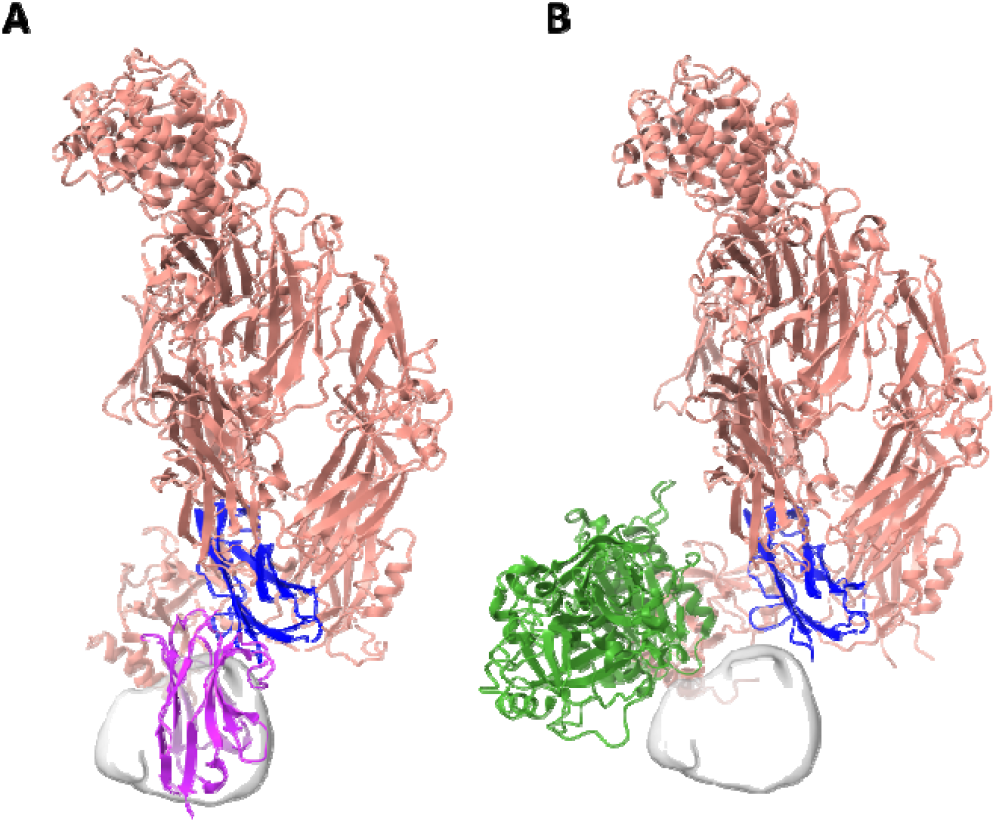
Two structures of C3 interacting with proteins via the MG7 domain **A)** Native C3 model (PDB 8OQ3) shown in red, MG7 domain highlighted in blue. The domain-specific nanobody (purple) is bound to the MG7 domain, inhibiting activation. The near identical location of TiO_2_ density (grey) from our current structure is shown for comparison. **B)** The C3 convertase model (PDB 2WIN) shown as red, MG7 domain highlighted in blue. The proximity of the factor B (green) and TiO_2_ density (grey) to MG7 show this area is vital for facilitating interactions with other regulatory complement proteins.

The MG7 domain plays a critical role in anchoring dynamic regions on the protein, such as the thioester domain, following C3 conformational changes. A recent report managed to deny the translocation of C3a and rearrangement of C3 by binding a specific nanobody to native C3.^41^ To achieve this, the nanobody was found to bind at the MG7 domain in the same position as the TiO_2_ nanoparticles. Therefore, the binding of the titania nanoparticles to MG7 has structural and physiological implications for the native motion of C3. The presence of the nanoparticles likely sterically hinders conformational shifts in C3, thereby locking the protein in an inactive state and disrupting intramolecular interactions necessary for the propagation of structural changes. This nanoparticle binding location demonstrates a charge complementarity between the highly electronegative, oxygen-coated, surface of the titanium nanoparticle and the charged, solvent-accessible residues on MG7. Although the resolution is not sufficient to unambiguously determine the conformation of the amino acids participating in the interaction, the location of the nanoparticle throughout all classification steps remained consistent, indicating this interaction as non-transient and stable. Importantly, no evidence from the structural or biophysical data suggests titanium nanoparticles are associated with C3 in the extended conformation or that they promote a major conformational shift to an active C3b-like state.

### Immune system implications of titania in the blood

The Compstatin family of cyclic peptides are inhibitors, specific for C3, and are shown to bind in a completely separate area of the protein. These inhibitor drugs are all documented to bind between the MG4 and MG5 domains, approximately 60 Å away from the observed TiO_2_ binding site at MG7. Yet, Compstatin binding does not arrest the conformation of C3, but instead prevents cleavage by convertases (C4b2a in the classical/lectin pathway and C3bBb in the alternative pathway). Large nanoparticles (>100 Å) are able to generate protein coronae, activate complement and initiate MAC formation in the blood. Given the biochemical and biophysical results all show a reduction in complement activity, it is likely that the TiO_2_ nanoparticles are recognised by the complement system as a substrate inhibitor of C3, rather than as a foreign surface.^42, 43^ Our findings uncover a previously uncharacterised mechanism by which TiO□ nanoparticles modulate innate immune responses via direct inhibition of complement component C3. Using nanoparticles that replicate the physicochemical properties - size, shape, and surface charge - of those known to leach from implanted medical and dental devices, we demonstrate selective suppression of the alternative complement pathway. This inhibition results in reduced generation of anaphylatoxins (C3a, C5a), impaired opsonisation, and decreased formation of the terminal membrane attack complex (MAC). Such activity suggests that TiO□ nanoparticles may alter host immune surveillance in the context of biomaterials, with implications for long-term implant biocompatibility and immune homeostasis. These results expand the functional profile of TiO□ beyond its established roles in structural support and surface engineering, reframing it as an active modulator of immune pathways. Our observations also align with growing evidence that metal and metal oxide nanoparticles—including silver, zinc oxide, cerium oxide, and gold—exert immunoregulatory effects through distinct molecular targets, warranting broader investigation into nanoparticle-host interactions in clinical materials.^44-46^ Efforts to further elucidate these mechanisms and assess their relevance in-vivo are ongoing in our laboratory.

## Materials and Methods

### C3 isolation, purification and storage

Native C3 was isolated and purified in-house from human plasma according to previous reports.^47^ The final step of purification involved cation exchange chromatography which was performed using a Mono S 5/50 GL column (GE Healthcare, Bio-Sciences AB, Uppsala, Sweden). The column was equilibrated with 5 column volumes (CV) buffer A (20 mM phosphate buffer pH 6.8), followed by sample application (0.25 CV). The column was washed with 3 CV of buffer A and then the elution was started using a linear gradient of buffer B (20 mM phosphate buffer pH 6.8, ranging from 0 to 0.85M NaCl). The protein elution was monitored with an UV detector at 280 nm and fractions of 0.5 mL were collected during the elution procedure. C3 was precipitated in a low salt and pH buffer for storage by dialysis against a 40 mM phosphate buffer with ionic strength 0.05 (mS) and pH 6.0 at +4°C with a buffer volume at least 10-fold higher than the volume of C3. After 24 h the buffer was replaced with fresh buffer and dialysis continued for another 24 h until a visible precipitate was formed. The stock solution of precipitated native C3 was then suspended (without centrifugation) and aliquoted before storage at -80°C. In order to dissolve the precipitated C3, 1-part VB^++^ (5x stock solution) was added to 4-parts of precipitated C3 immediately before use.^48^

### Preparation of sol-gel titania

The synthesis of the uniform titania colloids used in this work was performed by following a previously reported preparation.^49^ The precursor of titanium ethoxide (5 mL) was dissolved in anhydrous ethanol (5 mL) followed by the addition of 1.5 mL of triethanolamine during continuous stirring. A hydrolysing solution (1 mL) was produced by mixing nitric acid (0.5 M) with ethanol (2 mL). The resulting lightly-yellow, transparent solution contained 120 □ mg/mL TiO_2_ according to TGA measurements.^16^ The obtained product was of high-enough purity to not require additional purification.

### Biophysical measurements

DLS, thermal unfolding and nanoDSF measurements were performed on Prometheus Panta (NanoTemper, Munich, Germany) in technical triplicates. For apo-protein experiments, 0.5 mg/mL of C3 in filtered MQ water or buffered solution was added to separate capillaries at room temperature. For TiO_2_ experiments, nanoparticles in a stock solution of 1 mM were suspended in filtered MQ water, briefly incubated with C3 and added to capillaries. Immobilisation was carried out using a Biacore T200 system (GE Healthcare) at 25 °C with an AU bare gold, non-derivatised biosensor chip (Xantec) in Milli-Q water containing 0.05% Tween-20. A post-sonicated TiO □ solution at a concentration of 1.0 µg/mL was employed for immobilisation by simple manual deposition (10 uL/min), exploiting the strong affinity between the gold surface and titanium oxide. The immobilisation level was maintained at 300-400 RU. A reference channel was left untreated and did not undergo nanoparticle immobilisation.

Single-cycle kinetic experiments were performed with C3 in Tris buffer (50 mM Tris-HCl, 20 mM NaCl, 0.05% Tween-20, pH 7.4) under the following conditions: flow rate 75 µL/min, association time 60 s, dissociation time 900 s, and C3 concentration series prepared by two-fold dilution starting from 10 µM. A negative control with BSA confirmed the selectivity of the C3-TiO □ interaction, showing no detectable binding between BSA and the immobilised nanoparticles. Data analysis was performed using the Biacore Evaluation software. All raw data were exported for analysis and graph generation in GraphPad Prism version 10.0.

### Biochemical assays

For capillary electrophoresis analysis by Wes immunoassay and alternative pathway haemolysis, native C3 (1 µM) was preincubated with varying micromolar concentrations of TiO_2_ nanoparticles in VBS^++^ for 30 minutes at 37°C. For capillary electrophoresis, 70 µg/mL of C3, C3b and C3 incubated with nanoparticles were added to a preparation of Factor B (100 µg/mL) (Complement Technology Inc) and Factor D (0.5 µg/mL) (Complement Technology Inc) in VBS^++^ and incubated over a time course of 5, 30 and 60 mins. Individual time points were stopped with the addition of EDTA to a final concentration of 10 mM. The Wes immunoassay was performed under reducing conditions with a primary mouse monoclonal anti-human complement Factor Bb antibody (2 µg/mL) (Bio-Rad Laboratories).

The alternative pathway haemolysis assay was set up and adapted according to previous methods,^50, 51^ where 25 µL of C3 (200 µg/mL) preincubated with 0.5, 1, 2, 4, 8 and 16 µM of TiO_2_ nanoparticles were added to 50 µL C3 depleted serum (Complement Technology Inc) followed by the addition of 100 µL 50% rabbit erythrocytes (v/v) in GVB Mg EGTA. These samples were then incubated in a shaking water bath for 20 mins at 37°C at 200 rpm before being stopped with the addition of cold VB-EDTA. The samples were then centrifuged at 860 g for 10 mins at 4°C and absorbance measured at 540 nm on a CLARIOstar Plus (BMG Labtech).

### Cryo-EM structure determination of native C3

C3 protein buffer was exchanged from VB^++^ to filtered MQ water and concentrated to 0.4 mg/mL before grid freezing. Cryo-EM samples were prepared on UltrAuFoil R1.2/1.3 300 mesh holey gold grids (Quantifoil Microtools). Once the grids were glow discharged with a PELCO easiGlow instrument (20 mA, 0.4 mBar, 1.5 min), 3 µL of C3 was applied to the glow-discharged grids. Vitrification was performed with a Vitrobot Mark IV (Thermo Fisher Scientific) at 4 °C and 95 % humidity. Grids were blotted in duplicate for 3 seconds and plunged into liquid ethane. Following vitrification, the grids were clipped into autogrid cartridges (Thermo Fisher Scientific) for use with autoloader systems.

All grids were ATLAS screened on a Glacios 200 kV cryo-electron microscope mounted with a Falcon III direct electron detector. Data were acquired in EER format at a pixel size of 0.7463 Å/pixel with a dose rate of 0.91 electrons pixel^-1^ sec^-1^. Approximately 1200 micrographs had 2x binning applied and were processed using exposures fractionated into 50 frames for motion correction. This enabled for a rapid assessment of the sample quality, particle density and distribution before high-resolution data were collected. For full data collections, a 300 kV Titan Krios G2 (Thermo Scientific) transmission electron microscope equipped with a K3 detector (Gatan) and a 20 eV BioQuantum energy filter (Gatan), was used. Data were collected with the following settings: 0.648 Å/pixel (190 kx magnification), 51 e^−^/Å^2^ total electron dose, −0.5 to −2.5 μm defocus range. Automated collection was performed using the EPU software (Thermo Scientific) with 30° stage tilting. A total of 17,420 movies were collected.

During data collection, cryoSPARC live (cryoSPARC v4.1.1) was employed for on-the-fly data processing for the first few hundred micrographs using the same settings as for the initial screening dataset. Processing was performed in cryoSPARC v4.6.0^52^ and included initial Patch Motion Correction and Patch CTF being performed before micrograph curation. Micrographs were manually curated based on CFT estimations, total frame motion, defocus, ice thickness and contamination. Using the obtained template from the screening dataset, all particle picking was performed with template picker and the Topaz^53^ wrapper template picker in cryoSPARC. Once data collection was completed, the template-based picks totalled ∼5 million and were 2x downsampled and extracted with a 400 x 400 box size before being Fourier-cropped to 250 x 250 pixels. Particles were classified into 150 2D classes (10 Å minimum separation distance, 170 - 200 Å circular mask inner-diameter, 2 final full iterations, 40 online-EM iterations, 400 batch size per class). Following 2D class selection, this classification/selection procedure was repeated twice more which resulted in ∼1.9 million particles suitable for *Ab-initio* reconstruction. *Ab-initio* reconstruction was followed by heterogenous refinement using a few poor density “dummy” classes as seeds for junk particles. Two further rounds of heterogenous refinement seeding resulted in 818,343 particles which displayed discrete heterogeneity, which was elucidated using 3D variability analysis.^54^ Masked refinement and reconstruction jobs reliably produced densities at resolutions of ∼2.5 Å. 3D variability analysis demonstrated the inherent continuous disorder of the particles and was used to separate individual latent dimensions of motion attributed to approximately 3 states experienced by the particles. Particles responsible for each state were isolated and reconstructed independently to produce each individual C3 pose.

A model was constructed by using the reported inactive C3 structure (PDB 2A73)^36^ as a starting model. Owing to the large conformational differences between the current particles and the reported structure, the domains were individually docked and fitted into the electron density with UCSF ChimeraX version 1.9.^55^ These hand-fitted domains were then quantitatively fitted into the density and refined using the DockEM^56^ and Refmac Servalcat^57^ packages within CCP-EM 1.6.0.^58^ The Coot^59^ program was used for manual rebuilding and inspection of the model in the electron density before a final real-space refinement run in PHENIX.^60^ The relevant structural validation metrics are shown in supplementary Table 1.

### Cryo-EM structure determination of C3-TiO_2_

As previous, C3 protein buffer was exchanged from VB^++^ to filtered MQ water and concentrated to 0.4 mg/mL before grid freezing. Cryo-EM samples were prepared on UltrAuFoil R1.2/1.3 300 mesh holey gold grids (Quantifoil Microtools). While the grids were glow discharged with a PELCO easiGlow instrument (20 mA, 0.4 mBar, 1.5 min), C3 was incubated with nanoparticles at an equimolar concentration for 5 minutes on ice before 3 µL of the mixture was applied to the glow-discharged grids. Vitrification was performed with a Vitrobot Mark IV (Thermo Fisher Scientific) at 4 °C and 95 % humidity. Grids were blotted in duplicate for 3 seconds and plunged into liquid ethane. Following vitrification, the grids were clipped into autogrid cartridges (Thermo Fisher Scientific) for use with autoloader systems.

A 300 kV Titan Krios G2 (Thermo Scientific) transmission electron microscope equipped with a K3 detector (Gatan) and a 20 eV BioQuantum energy filter (Gatan), was used for the full data collections. Data were collected with the following settings: 0.648 Å/pixel (190 kx magnification), 51 e^−^/Å^2^ total electron dose, −0.5 to −2.5 μm defocus range. Automated collection was performed using the EPU software (Thermo Scientific) and a total of 17,706 movies were collected. Processing was performed in cryoSPARC v4.6.0 and included initial Patch Motion Correction and Patch CTF being performed before micrograph curation. Micrographs were manually curated based on CFT estimations, total frame motion, defocus, ice thickness and contamination.

A visual inspection of the micrographs and blob-picked classes in cryoSPARC suggested there was a clear picking bias which made further classification and alignment impossible (ESI Figure S4a). As the electron dosage charged the TiO_2_, the nanoparticles appeared as bright hotspots on the micrographs which almost completely overpowered signals attributed to protein. Attempts to use the previous C3 reconstruction as a picking template were unsuccessful due to the picking algorithm being unable to locate any signal related to protein. Analysis of the protein and TiO_2_ relative intensities in EMAN2^61^ showed that there was approximately a 1:10 intensity disparity between the protein and nanoparticles, respectively (ESI Figure S4c). Attempts to enforce non-negativity, solvent-clamping and template picking using native C3 prior to 2D classification proved unsuccessful as the picking was strongly biased toward the most intense signals of the nanoparticles. To the best of our knowledge, cryoSPARC does not have the ability for the user to manually threshold and normalise intensities, and for this reason we attempted to complete the data processing using RELION 5.0.^62^ Following a standard pre-processing strategy up to CTF estimation, Topaz was trained on a small subset of particles and once training converged, particles were extracted from all micrographs. The extraction job settings were not default: particle box size 384 pixels, diameter background circle 40 pixels, stddev white dust removal -1.0, stddev black dust removal 3.0, with normalisation on and a minimum FOM of 1.0. The relion image handler tool was used to threshold a series of incremental values from 1.0 – 3.0 in order to find which cutoff yielded the best SNR of both protein and nanoparticle. After 2D classification was run on each thresholded particle stack, normalisation was run using the relion stack create and relion preprocess tools. Performing this step was found to improve particle alignment as well as provide an intensity distribution favoured by RELION. A low-resolution map was obtained from the thresholded and normalised particle stack after running through 2D classification, 3D reconstruction and masked refinement.

The obtained density showed most domains present excluding those in the macroglobulin head region (MG1-4). Excluding this missing area, the domains were individually docked and fitted into the electron density with UCSF ChimeraX version 1.9, which conformed to the inactive state C3 conformation.

### Structural and statistical image preparation

All cryo-EM images were prepared using ChimeraX. All biophysical and biochemical graphs and analyses have been prepared, fitted and displayed using the program GraphPad Prism version 10.0.

## Acknowledgements and funding

We gratefully acknowledge the Cryo-EM Uppsala facility for the use of the Glacios microscope, computational facilities and the expert technical support of Daniel Larsson, all of which is funded by the Department of Cell and Molecular Biology, the Disciplinary Domains of Science and Technology and of Medicine and Pharmacy at Uppsala University. This work was supported by the SciLifeLab & Wallenberg Data Driven Life Science Program, Knut and Alice Wallenberg Foundation (grants: KAW 2020.0239 and KAW 2017.0003), and by the National Bioinformatics Infrastructure Sweden (NBIS) at SciLifeLab. We would like to especially thank Dustin Morado, Tim Schulte, Piotr Draczkowski and Helena Danielson for their expert assistance with data collection and interpretation. The computations and data handling were additionally enabled by resources provided by the National Academic Infrastructure for Supercomputing in Sweden (NAISS), partially funded by the Swedish Research Council through grant agreement no. 2022-06725. This work was supported by the Swedish research council grant number 2022-03971_VR. We kindly thank Guillermo Pérez Ropero and Ridgeview Instruments AB for the C1 SPR sensor chip donation.

## Author contributions

M.S, K.N.E, B.N and V.G.K conceptualised the research. G.A.S synthesised the pure titania colloidal nanoparticles. J.W performed SPR, data analysis and generated the figures and text. D.E and K.F purified the C3 protein and performed all biochemical and haemolysis assays. J.J.W performed DLS and nanoDSF characterisations with C3 and TiO_2_. J.J.W prepared all TEM grids, performed data collection, processing, model refinement and validation of all cryo-EM structures. J.J.W and D.E compiled and analysed the data, wrote the original manuscript and prepared figures and tables. Editorial assistance was provided by V.G.K, G.A.S, M.S, B.N and K.N.E. All authors have reviewed, edited and approved the final version of the manuscript.

## Data availability

Original data points are shown in the manuscript. Additional information can be obtained from the corresponding author upon request.

