## Supplemental Information for "Titanium Nanoparticle Regulates Innate Immunity"

Electronic Supporting Information

List of supplementary table and figure captions

**Supplementary Table 1.** Cryo-EM data collection, refinement and validation statistics for native compact C3, stretched C3 and C3-TiO_2_.

**Supplementary Figure S1.** SDS-PAGE of purified complement C3 following size exclusion chromatography. All bands can be assigned exclusively to native C3 while bands for C3b were unable to be observed.

**Supplementary Figure S2.** 3D variability component details of native C3. **A)** A pair-wise comparison of electron density shifts from a consensus map between two components. Red regions show an increase in electron density within the coloured area, while blue represents a decrease in density. **B)** A heat map showing the distribution of density over a latent dimension. The two supplemented circles show two dark red “hot spot” zones, suggesting the presence of two distinct conformational components within the data.

**Supplementary Figure S3.** DLS and nanoDSF investigations into the relative stabilities of native C3, TATT nanoparticles and C3-TiO_2_. **A)** Technical duplicate isothermal DLS experiments of TiO_2_ nanoparticle in buffer (Tris 50 mM, NaCl 50 mM, pH 7.4). Immediate particle aggregation is evident from timepoint zero through to 90 minutes. **B)** The repeated isothermal DLS experiment of TiO_2_ in filtered water at a range of concentrations. All conditions showed the expected nanoparticle size and remained stable over 2 hours. The elimination of salt and buffered conditions significantly improves the homogeneity and stability of the nanoparticles. **C)** NanoDSF experiments showing the increased stability of C3 in the presence of 100 µM or 500 µM of TiO_2_ in both filtered water and buffer. An increase in Tm from 59 $℃$ (red) to 66 $℃$ (blue) was found for native C3 with 100 µM of TiO_2_ in filtered water.

**Supplementary Figure S4.** 2D classifications of C3-TiO_2_ showing the before and after using cryoSPARC and RELION-5 pipelines. **A)** Post-particle picking and extraction in cryoSPARC produced 2D classes consisting of bright spots from the charged nanoparticles. **B)** The 2D classes shown in RELION-5 are the result of manual normalisation and signal thresholding during extraction, allowing for an attenuation between the imbalanced signals, increase in SNR and a greatly-improved particle alignment. **C)** The 10-fold relative intensity factor discrepancy between the inorganic nanoparticles and organic protein species. **D)** Zoomed-in classes showing (blue) inactive conformations of native C3 with surrounding black spheres of nanoparticle. (Orange) Selected TiO_2_ nanoparticle classes – each black sphere is ~3.5 nm in diameter.

**Supplementary Figure S5.** NanoDSF experiment of native Factor B and Factor B–TiO_2_. The black curve shows 0.5 mg/mL of apo-Factor B in filtered water with a corresponding Tm of ~55 $℃$. Technical duplicates of factor B in the presence of 100 µM of TiO_2_ produced a near-identical Tm with no apparent adverse or beneficial effect on protein stability, suggesting no meaningful interaction exists between the two species.

**Supplementary Figure S6.** Wes immunoassay, reducing conditions, size range of 12-230 kDa. Generated from Compass for Simple Western software showing C3, C3b and C3 TATT incubated samples detected with a mouse anti-human Factor Bb neoepitope. Three experiments incubating Factor B and Factor D with one of the following C3 preparations. C3, C3b (positive control), C3 pre-incubated with nanoparticles (incubated for 5, 30 and 60 mins).

**Supplementary Figure S7.** 2D classes of native C3 particles with a corresponding GS-FSC curve following refinement. **A)** 50 2D classes are shown which demonstrate the variety of conformations possible from native C3 in filtered water. Notably, no conformations suggest the native C3 has been activated to a C3b conformation. **B)** Gold-standard Fourier Shell Correlation (GS-FSC) curves are shown, indicating a resolution of 2.58 Å at FSC_0.143_.

**Supplementary Table 1. Cryo-EM validation table**

| Molecule | Native C3 | Native C3 extended | C3-TiO_2_ |
| --- | --- | --- | --- |
| Data and processing |  |  |  |
| Magnification | x190,000 | x190,000 | x190,000 |
| Voltage (kV) | 300 | 300 | 300 |
| Defocus range (µm) | − 0.6, −2.0 | − 0.6, −2.0 | − 0.6, −2.0 |
| Pixel size ($\boldsymbol{Å}$) | 0.648 | 0.648 | 0.648 |
| Symmetry | C1 | C1 | C1 |
| Stage tilt (°) | 30 | 30 | 0 |
| Electron exposure (e^-^/ $\boldsymbol{Å}$^2^) | 51.1 | 51.1 | 51.7 |
| Initial particle images (no.) | 3,630,401 | 729,181 | 345,569 |
| Final particle images (no.) | 408,361 | 101,194 | 84,518 |
| Map resolution at FSC_0.143_ ($\boldsymbol{Å}$) | 2.58 | 4.1 | 7.1 |
| EMDB code | EMD-53900 | EMD-53939 | EMD-53940 |
| PDB code | 9RBO | 9RF3 |  |
| Refinement |  |  |  |
| Model resolution ($\mathbf{Å}$) | 3.0 | 5.8 |  |
| FSC threshold | 0.143 | 0.143 |  |
| Map sharpening | -83.0 | -48.0 |  |
| *B factor (*$\boldsymbol{Å}$*^2^)* |  |  |  |
| Atoms (non-H) | 11,688 | 11,570 |  |
| Residues/ligands | 1,474 | 1,459 |  |
| *ADP B factors (*$\boldsymbol{Å}$*^2^)* |  |  |  |
| Protein/ligands (min/max) | 33.04/174.54 | 67.61/153.44 |  |
| *R.m.s deviations bonds* |  |  |  |
| Lengths ($\boldsymbol{Å}$)/angles (°) | 0.005/1.178 | 0.004/0.853 |  |
| Validation |  |  |  |
| MolProbity score | 3.02 | 2.72 |  |
| Clash score | 28.49 | 27.40 |  |
| Poor rotamers (%) | 1.52 | 1.86 |  |
| *Ramachandran plot* |  |  |  |
| Favoured | 81.92 | 81.53 |  |
| Allowed | 16.78 | 18.13 |  |
| Disallowed (%) | 1.30 | 0.34 |  |

**Supplementary Figure S1.**

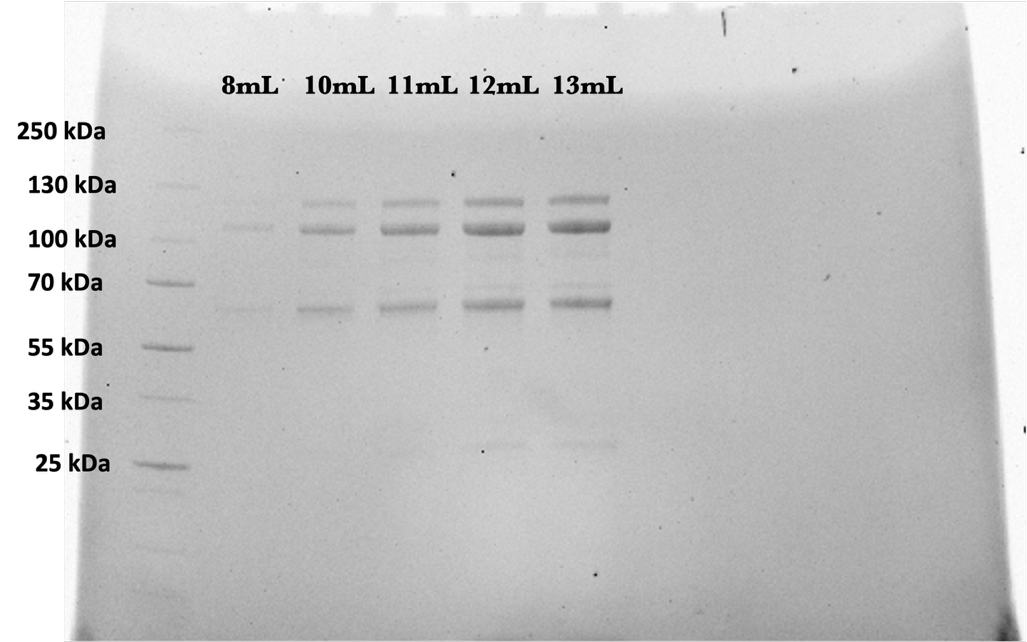

**Supplementary Figure S2.**

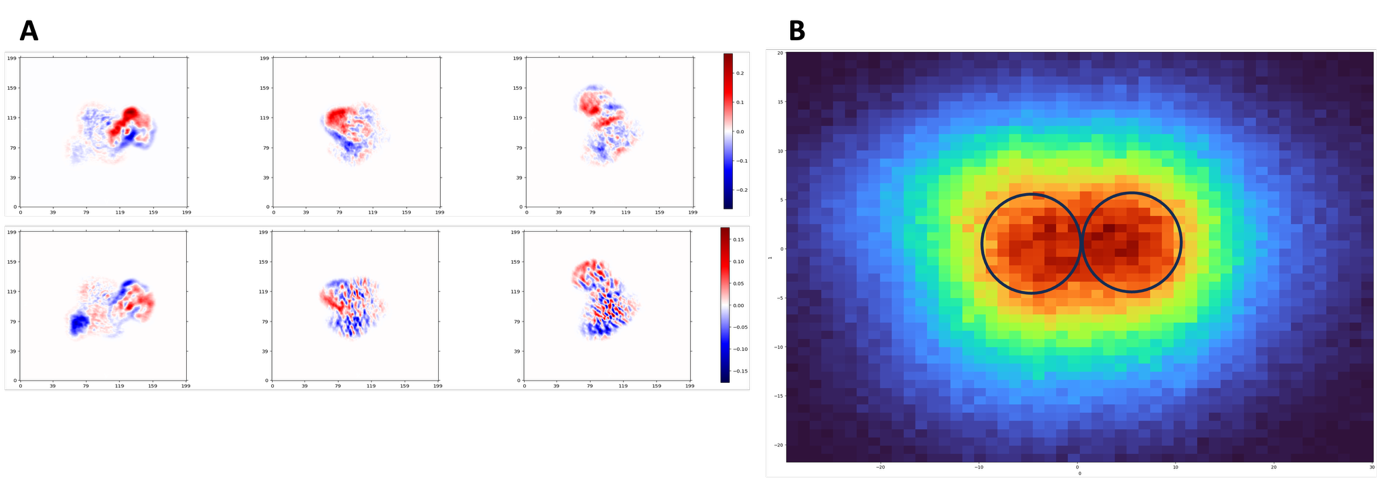

**Supplementary Figure S3.**

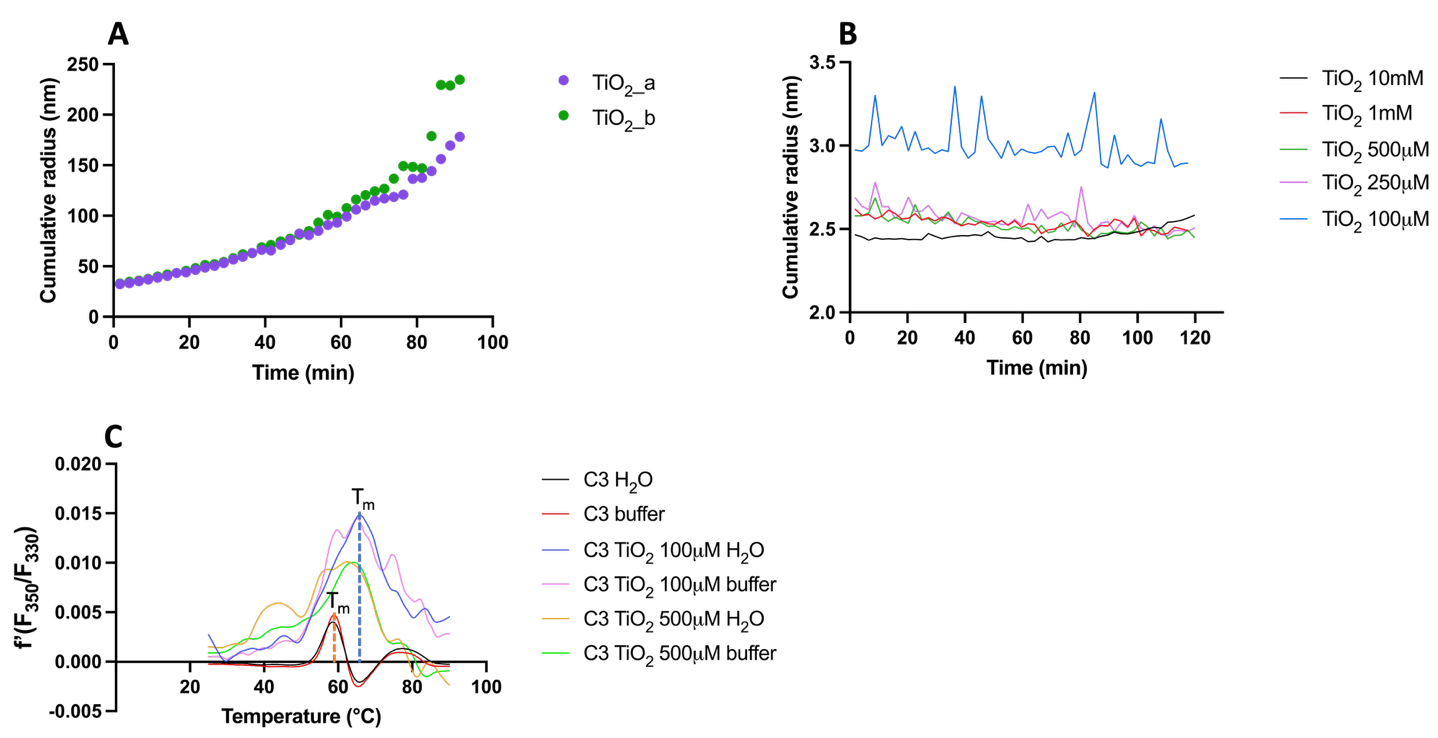

Biophysical analyses of complement C3

Prior to structural analyses, baseline characterisations of native C3 were undertaken, including melting temperature (T_m_) and hydrodynamic radius (R_h_) at physiological and extreme conditions. DLS experiments showed that within the range of 25 – 55 $^{\circ}$C, the protein maintained a constant diameter of ~12 nm (120 Å) which, surprisingly, is 40 Å smaller than reported inactive C3 structures (Figure 1a).^28, 29^ Polydispersity values were consistently low and the hydrodynamic size remained monodispersed and unchanged after 90 minutes. Thermal stability tests demonstrated that heating the sample incrementally produced notable and rapid C3 aggregation at around 58 $^{\circ}$C, with 6 nm cumulative radii protein reaching 100 nm at 62 $^{\circ}$C. This result was supported by nanoDSF measurements which showed fast protein denaturation and a resulting T_m_ of 60 $^{\circ}$C (Figure 1b).

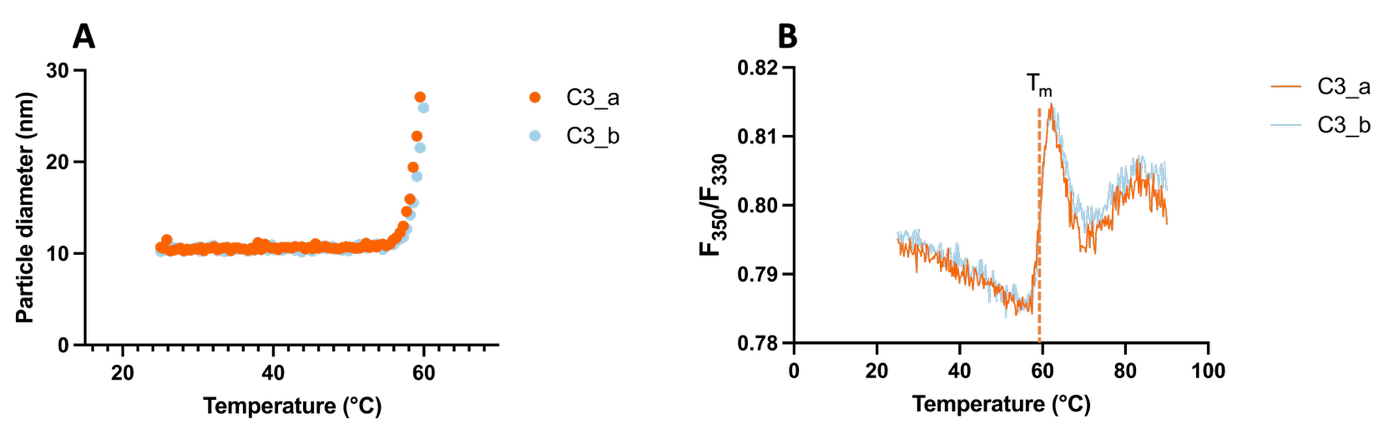

**Figure 1.** DLS and nanoDSF investigations into the relative stabilities of native C3. **A)** Overlaid DLS experiments of C3 at 0.4 mg/mL concentration in buffer (Tris 50 mM, NaCl 50 mM, pH 7.4, orange) and in filtered water (blue). **B)** Overlaid nanoDSF runs of 0.4 mg/mL native C3 in buffer (Tris 50 mM, NaCl 50 mM, pH 7.4, orange) and native C3 in filtered water (blue). Both melting and particle aggregation temperatures are near identical, represented by the dashed line at the inflection point (Tm $\approx$ 59 $℃$).

SDS-PAGE and size-exclusion chromatography were used to confirm whether the smaller-than-expected protein diameter was due to missing or cleaved domains besides the 9 kDa anaphylatoxin domain of C3a. Despite possessing the same number of domains, the inactive and hydrolysed states of complement C3 have been shown to elute at significantly different times simply owing to considerable inter-domain reorientation.^30-32^ Following protein digestion under reducing conditions, the gel showed all bands and their corresponding domains were accounted for. The size-exclusion chromatogram further confirmed that all domains were present with the 2 separate elution peaks being attributed to both inactive and hydrolysed C3. Importantly, no smaller peaks or contaminants were detectable after the final C3-associated elution (ESI Figure S1).

Although all biophysical and purity analyses showed the C3 sample as homogenous, conflicting evidence suggested the protein was significantly smaller than the crystallographic and cryo-EM structures already reported. To explain this discrepancy, we turned to single-particle cryo-EM and determined the structure of native complement C3 under the same physiological conditions as all previous analyses. To our surprise, we observed several states of the complement protein achieving an extreme range of inter-domain motion.

**Supplementary Figure S4.**

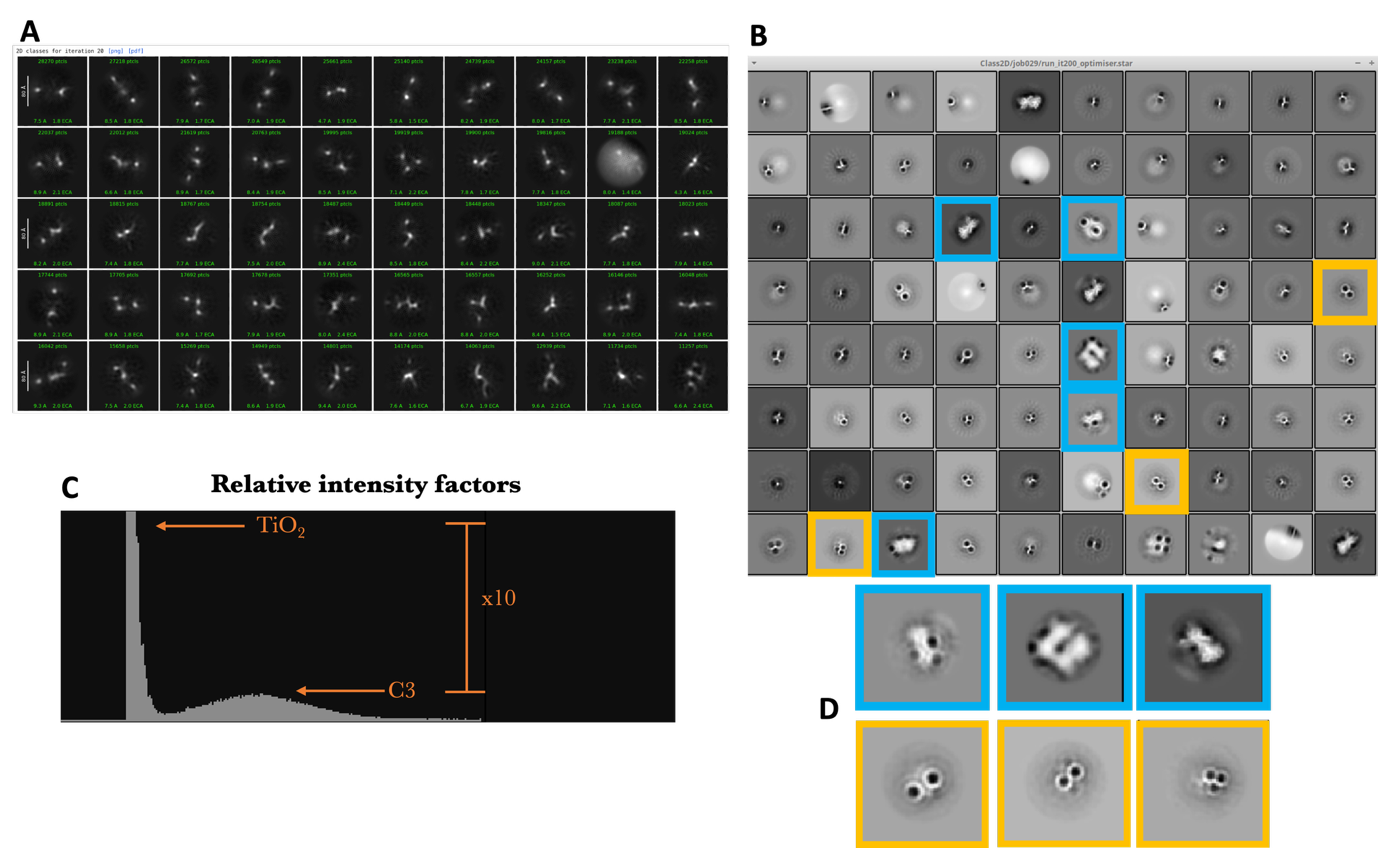

**Supplementary Figure S5.**

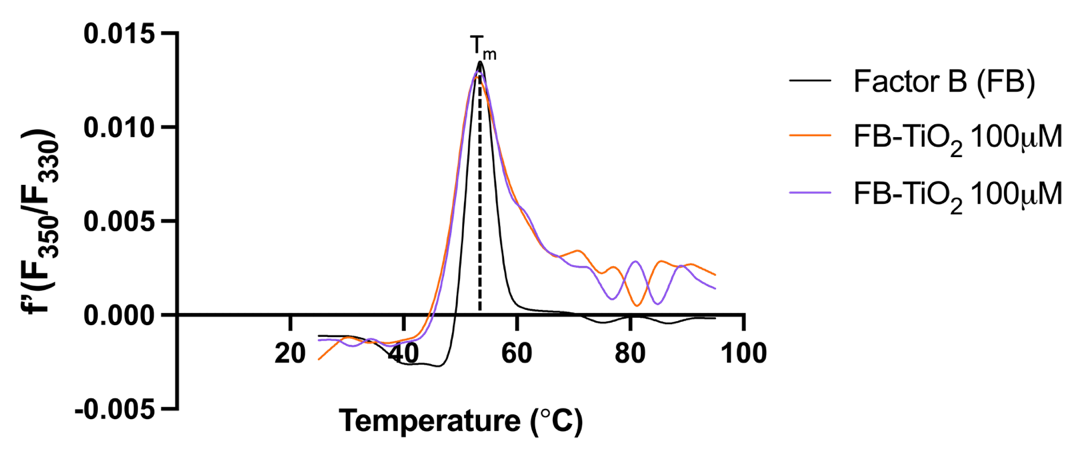

**Supplementary Figure S6.**

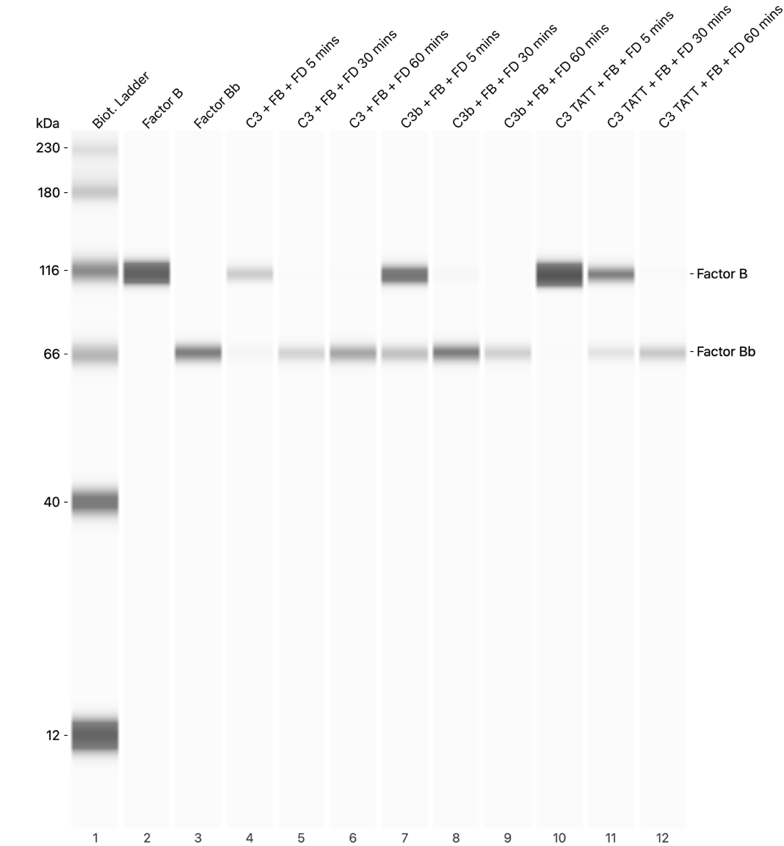

**Supplementary Figure S7.**

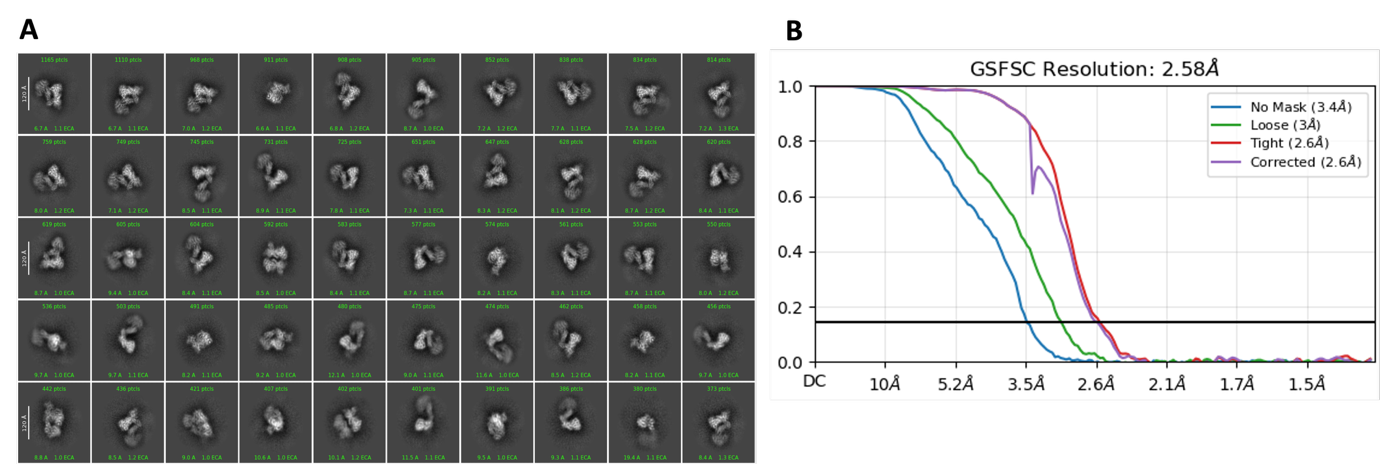
